# Do heatwaves impact mosquitoes? Multi-level analyses across developmental stages in *Aedes albopictus*

**DOI:** 10.1101/2025.07.01.662530

**Authors:** Claudia Alfaro, Alejandro Nabor Lozada-Chávez, Ayda Khorramnejad, Alida Kropf, Paolo Luigi Catapano, Alessia Cappelli, Claudia Damiani, Guido Favia, Mariangela Bonizzoni

## Abstract

The increase in the frequency and intensity of heatwaves (HWs) under global warming is expected to have more dramatic biological impacts on insects than mean temperature increases by challenging their thermal thresholds. However, experimental evidence that quantifies the impact of HWs on insects in their ecological context is limited. Here, we measured across multiple biological scales (e.g. fitness, physiology, transcriptomic and microbiota) the stage-specific responses of the arboviral vector *Aedes albopictus* when experiencing an ecologically relevant HW. In arboviral vectors, only females require a blood-meal thus contributing to the transmission cycle, but males, eggs and larvae can also be the target of control strategies to reduce population size. As such the understanding of the responses to a HW during mosquito development has both biological and epidemiological relevance.

We observed stage-specific responses, starting at the egg stage. We saw egg hatching decreasing and delaying after HW exposure. On the contrary, larvae showed to be resilient to HW. Larvae repurposed their energy resulting in trivial mortality, but delayed development. At the adult stage, we observed a marked sex difference, with extensive (>50%) male mortality, accompanied by a small (16) number of genes elicited following HW exposure, which indicates that males have limited coping mechanisms against hot events. In females, we observed a reduced reproductive output, but only when HW occurred after a blood-meal. Finally, we saw extensive HW-dependent changes in the microbial composition of larvae, but female microbiota remained dominated by *Wolbachia* regardless of the thermal challenge. Our results have relevant implications for both the understanding of mosquito biology and the implementation of vector control strategies as the climate crisis progresses.

## 1. INTRODUCTION

Being dependent on environmental temperature (Ta) to modulate their physiology and behaviour, insects are particularly susceptible to variation in Ta, including both gradual Ta increase and abrupt Ta shifts to extreme values, broadly defined as coldsnaps or heatwaves (HWs), depending on the direction of the thermal variation with respect to average Ta (Mirth et al., 2021). Under current global warming, HWs are more prevalent than coldsnaps and increasing in frequency (Meehl & Tebaldi, 2004). HW exposure is known to exert profound effects on insects by activating stress response genes (*i.e.* heat shock proteins, detoxification enzymes and protein kinases) and eliciting thermal tolerance mechanisms, which are energy demanding (Clissold & Simpson, 2015; Colinet et al., 2015; González-Tokman et al., 2020; Hendrix & Salvucci, 1998; Lahondère, 2023; Nguyen et al., 2009). However, the magnitude of these responses, and consequent fitness costs, are highly variable. They depend not only on HW characteristics, including the level of the thermal shift, its frequency and fluctuating regime, but also on the insect species and its developmental stages (Walsh et al., 2019). Different instars may have distinct thermal sensitivities, resulting in stage-specific thermal responses (Kingsolver et al., 2011; Pincebourde & Casas, 2015; Ren et al., 2023). As a result, the fitness costs to a HW of the different life-stages of an insect may vary, also in relation to their environment (i.e. aquatic or terrestrial), the accessibility of, and possibility to, move to alternative microclimates, the size of their body and their metabolic state (Kingsolver et al., 2011; Sales et al., 2021; F. Zhao et al., 2017). At present, responses to a HW have been quantified in a limited number of model insect species and often not thoroughly across biological scales, such as fitness, physiology, transcriptome thus hindering a comprehensive understanding of the organism’s rebound to a hot event (Li & Gong, 2017; Zani et al., 2005) and challenging our abilities to predict the impact of future HWs on insects in different ecological contexts. In mosquito species, which can vector pathogens to humans, assessing the impact of a HW across developmental stages and sexes has both ecological and epidemiological relevance. Only females contribute to the transmission cycle based on their dependency on blood-meals from vertebrate hosts, but control strategies to reduce population size may also target males or eggs and juvenile stages, which occupy different ecological niches than adults (Achee et al., 2019).

To probe the biological impacts of a HW on mosquitoes, we chose the Asian tiger mosquito *Aedes albopictus*, which is a highly invasive species and the primary vector of arboviruses in temperate areas of the world (Bonizzoni et al., 2013). Originally from Southeast Asia, *Ae. albopictus* established in Europe and North America in the early 1990, leading to the emergence of autochthonous arboviral outbreaks (Aranda et al., 2018; Bohers et al., 2024; Farooq et al., 2025). In a previous study, we showed that the thermal optimum for *Ae. albopictus* ranges from 23°C to 26°C depending on the geographical origin of the tested populations (Carlassara et al., 2024). Building on this result, we exposed the same laboratory populations to a HW as recorded in the summer of 2021 in Rome (World-Weather, https://world-weather.info/forecast/italy/rome/august-2021/) and quantified life-history traits across mosquito development (Figure 1). We further explored the transcriptional responses of larvae and adults exposed to a HW, measured adult energy reserves to understand the phenotypic responses at a molecular level, and studied HW-dependent changes in mosquito microbiota. We observed that larvae are more resilient to a HW than adults and that males suffer more costs than females for which nutrients from a blood meal contribute to overcome HW-related protein depletion. Finally, we saw that while the microbiota of larvae was significantly alerted following HW exposure, that of females remained dominated by *Wolbachia*. Overall, our results have relevant implications for both the understanding of mosquito biology and the implementation of vector control strategies as the climate crises progress.

**Figure 1.**
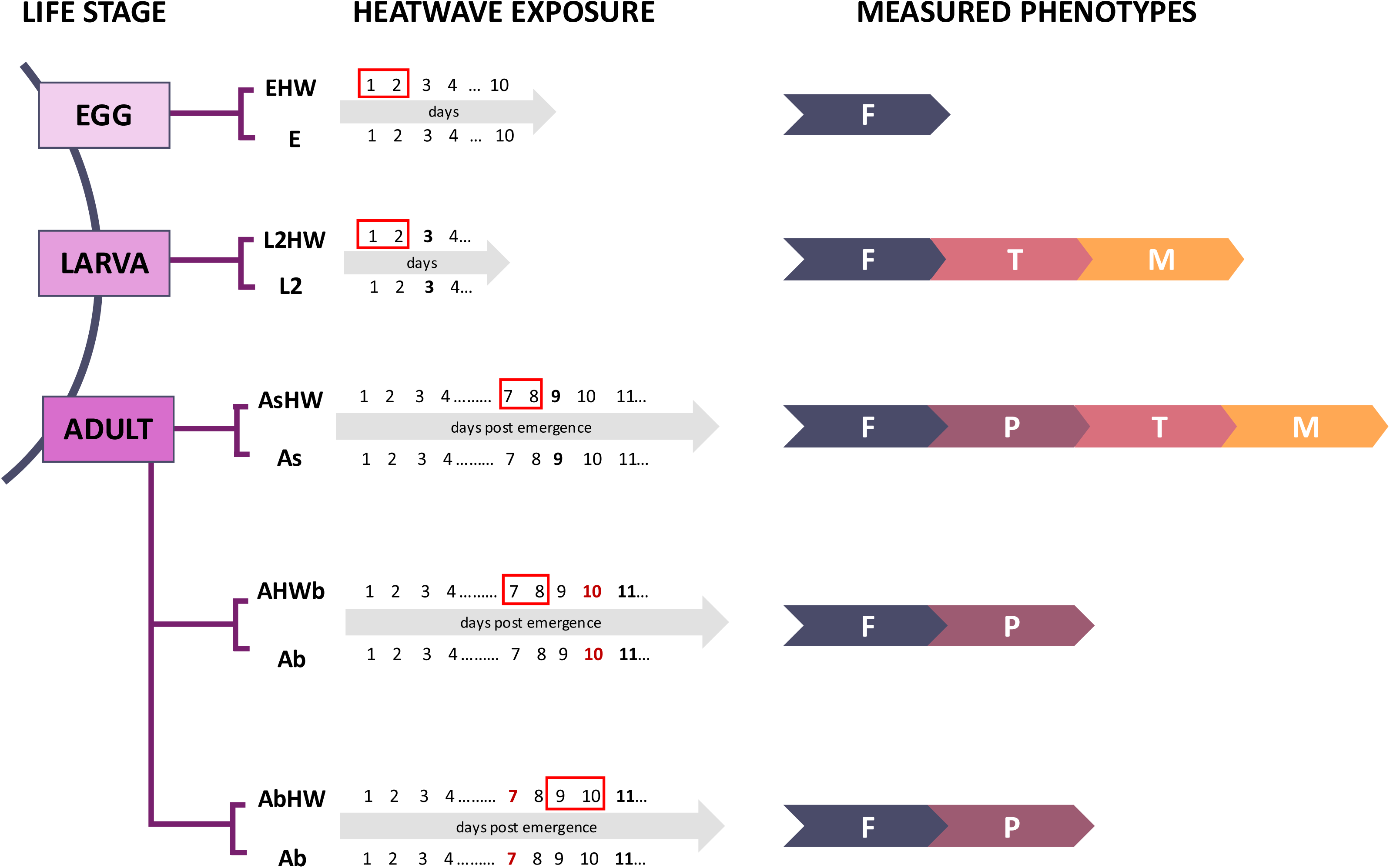
Experimental design. We exposed to a heatwave (HW, red box) consisting of 2 days at 37°C for 12 hours followed by 26°C for 12 hours eggs (E), larvae (L2), sugar-fed adults (As) and females offered a blood-meal either before (AbHW) or after (AHWb) the HW. Following the HW and/or the blood-meal, we measured fitness traits (F) and energy reserves (P). For L2 and As samples, we further analysed transcriptional (T) and microbiota (M) changes following HW. For each treatment, controls were done as indicated in the line below the arrow, which lists the days post emergence. The day in which the blood meal was offered is in red.

## 2. MATERIALS AND METHODS

### 2.1 Mosquitoes

We used *Ae. albopictus* of the genome-reference Foshan (Fo) strain (Palatini et al., 2020), along with mosquitoes adapted to the laboratory from eggs collected in the city of Crema (Cr) in Italy in 2017 (Carlassara et al., 2024). Since laboratory adaptation, both strains have been maintained in Binder KBWF climatic chambers under standard rearing conditions of 12h at 28°C and 12h at 26°C (light:dark), with 70% relative humidity. We rear larvae in BugDorm plastic containers (19×19×16 cm) at a density of 200 larvae for 1 L of water. Food is provided daily in the form of fish food (Tetra Goldfish Gold Colour). Adults are kept in 30 cm^3^ cages and fed with cotton soaked in 20% sugar solution. Each cage contains approximately 150 adults. Females are bloodfed with commercial defibrinated mutton blood (Biolife Italiana) using a Hemotek blood membrane feeding apparatus.

### 2.2 Heatwave

Using a Binder KBWF, we mimicked a HW that lasted two days and consisted of a cycle of 12h at 37°C, followed by 12h at 26°C (light:dark, 70% relative humidity) (Figure S1a). After HW exposure, mosquitoes were moved back to standard rearing conditions.

### 2.3 Assessment of fitness traits

For each strain, we measured: hatching rate, larval viability and larval developmental time, pupae viability and pupae developmental time, female and male longevity of virgin mosquitoes, and their reproductive capacity. We describe below how each fitness parameter was measured.

#### Hatching rate

For each strain, three groups of 350 eggs were hatched in plastic containers (19×19×16 cm) with 1L of water at 37°C and immediately placed into the Binder KBWF set to the HW regime. After the HW, trays were moved to standard rearing conditions. In parallel, three groups of 350 eggs were hatched in 1L of water at 28°C and kept under standard rearing conditions. For each tray, we counted larvae that hatched daily for ten days. The percentage of emerging larvae with respect to the initial number of eggs was used as a measure of hatching rate.

#### Larvae and pupae fitness parameters

For both strains, we exposed a total of 120 second instar larvae (L2) to our HW. After HW exposure, we counted the number of dead larvae to assess heatwave-related mortality. We transferred survivors to the standard regime and tracked their growth daily to measure larval developmental time (LDT) as the number of days it took until pupation, larval viability (LV) as the percentage of pupae with respect to initial larvae, pupal developmental time (PDT) as the time of adult emergence from pupae and pupal viability (PV). Finally, we calculated the sex ratio counting the number of females and males that emerged.

#### Adult fitness parameters

Experiments were conducted in parallel for both strains. For each strain, we exposed 5-7 days old mosquitoes to the HW, we exposed between 200 and 250 individuals of each sex/strain. Immediately after HW, we counted the number of dead adults to measure HW-related mortality and surviving adults were sex-sorted to follow females and male longevity separately. Longevity after HW exposure was assessed in a total of 240 individuals in groups of 80 individuals per replicate. Besides longevity we also checked the effect of the HW on the reproductive capacity of both males and females. For males, a total of 35 males/strain surviving HW exposure were allowed to mate with unexposed virgin females; we set up crosses following a ratio of 1 male:4 females in groups of 5 males per cage. After 7 days, females were offered a blood meal, and both fecundity and fertility were measured as described by Tsujimoto & Adelman, (2021). We further confirmed the mating status of females that had not deposited eggs by dissecting and viewing their spermatheca under an Olympus CKX53 microscope.

We tested the reproductive capacity of females, which were exposed to the HW either before and after the blood meal. In the former, females were allowed to recover from the HW for 24 hours before a blood meal was offered and we also counted the number of females acquiring a blood meal to check if blood feeding rate is affected by HW exposure. In the second scenario, that is exposure to HW after the blood meal, we offered a blood meal to 5-7 day old females and waited 24 hours before exposing them to the HW. For both scenarios, we had a group of control mosquitoes maintained under standard conditions. In all cases, fecundity and fertility was measured for 48-54 females for each strain under standard conditions.

### 2.4 Energy reserves quantification

We quantified protein, carbohydrate, glycogen and lipid content using a colorimetric assay as previously described (Carlassara et al., 2024) in 30 males and 30 sugar-fed females after HW exposure and their respective controls. Adults were 5-7 days old when they were exposed to the HW. We further quantified protein content in 30 females that had taken a blood meal followed by HW exposure and their respective controls. We weighed individuals using a microbalance (Mettler AC100). Then, using pestles we homogenized samples individually in 180 μL of lysis buffer (100 mM KH2PO4 (Sigma-Aldrich), 1 mM DTT, 1 mM EDTA (ThermoFisher) pH 7.4) and measured absorbance using a CLARIOstar plate reader (BMG Labtech).

### 2.5 Midgut trypsin-like activity

One day after females surviving the HW took a blood meal, we measured their midgut trypsin-like activity. On ice, we dissected midguts of at least 36 females per strain and tested trypsin enzymatic activity as previously done (Gulia-Nuss et al., 2012). Briefly, we used pestles to homogenize each sample in 100 µL extraction buffer (20 mM Tris/ 20 mM CaCl2 [Sigma-Aldrich], pH 8). Then, we centrifuged samples for two minutes at 14000*g* at 4°C, we collected the supernatants and stored them at –80°C until measurements. We added 5µL of each sample to 100µl of Nα Benzoyl-L-Arginine-p-Nitroanilide (BApNA; Sigma-Aldrich) in a 96-well plate, which were then incubated at 37°C for 10 minutes. We read the absorbance at 405nm using a CLARIOstar plate reader (BMG Labtech). We quantified trypsin-like activity using 20µg of trypsin from Bovine Serum (Sigma-Aldrich) for the standards.

### 2.6 Microscopy analyses of ovaries

From each strain, we collected 10 sugar-fed females and 10 females that had been offered a bloodmeal before HW exposure and we examined ovary resorption immediately after HW as described by Clifton & Noriega, (2012). Briefly, we dissected ovaries and rinsed them in physiological saline buffer (APS; MgCl2 0.6mM, KCl 4 mM, NaHCO3 1.8mM, NaCl 150mM, HEPES 25 mM, CaCl2 1.7mM [all Sigma-Aldrich]). Afterwards, we stained ovaries with 0.5% neutral red solution (Sigma-Aldrich) in acetate buffer pH 5.2 (Sigma-Aldrich) for 10 seconds, rinsed them again in APS and placed them under a coverslip. We visualized samples using an Olympus CKX53 microscope.

### 2.7 Transcriptome analyses

We collected three pools of 10 L2 larvae of the Cr strain immediately after HW and three pools of 10 L2 kept at standard rearing conditions as control. We also sampled four Cr females and four Cr males 24 hours before HW, and immediately after HW exposure; hereafter we will refer to these samples as As6 or As9HW because all samples were six or nine days old, respectively. We used these samples to test a potential ‘two-days time effect’ resulting from HW exposure (see Figure S1). We additionally collected the same number of females and males that had been kept at standard rearing conditions (i.e., control conditions) for the period at which the other experimental group had been exposed to the HW; hereafter we will refer to these samples as As9 as by the time of the collection they had reached nine days post emergence (Figure S1). We dissected the abdomens of each adult. Larval pools and individual abdomens were homogenized in 50µL of Trizol (Life Technologies) and stored at –80°C until extraction. We performed RNA extraction following the standard Trizol procedure and re-suspended samples in a final volume of 20µL of nuclease-free water. We sent total RNA to BGI Hong Kong for quality control, TruSeq-Stranded mRNA library preparation and sequencing on the Illumina platform. Libraries were sequenced pair-end (2×100 bp) at a depth of 30 million reads.

For RNAseq analysis we used the nf-core/RNAseq pipeline v3.17.0 (Patel et al., 2025) with the latest AalbF5 genome assembly of *Ae. albopictus* (RefSeq: GCF_035046485.1). The pipeline includes quality check of reads and their trimming before genome mapping for gene expression quantification. We included sequence reads corrections in our nf-core/RNAseq command line for salmon to prevent a “random hexamer priming bias” and a “GC bias”, as suggested (van Gurp et al 2013; Patro et al 2017). We used fragments per kilobase of exon per million mapped reads (FPKM) to quantify gene expression (Table S1). First, we defined a gene as expressed when the gene reads counts ≠ 0, which allowed us to estimate the total fraction of genes expressed under a HW across the AalbF5 whole genome for different life stages (Table S2). Overall, in the larval samples, we detected a total of 18151 expressed protein-coding genes in the control conditions, and a total of 18354 expressed protein coding genes in samples exposed to the HW, which accounts for 76.80% and 77.66%, respectively, of the protein coding genes of the *Ae. albopictus* genome (Table S2). In samples from adult females, we detected 18610 expressed protein-coding genes under control conditions and a total of 18376 expressed protein coding genes in samples that had been exposed to the HW, which represent 78.74% and 77.75%, respectively, of the protein-coding genes of the *Ae. albopictus* genome. Finally, in males we detected 18916 and 19090 expressed protein-coding genes in control samples and samples exposed to the HW, respectively. This represents 80.03% and 80.77% of the protein-coding genes of the *Ae. albopictus* genome for the former and later samples, respectively. Second, we performed a Principal Component Analysis (PCA) with RStudio v4.4.1 to show the association between samples based on the raw reads counts and developmental stage, sex and treatment (Figure S2a). Third, we compared gene expression across conditions and life stages using the DeSeq2 package v1.46.0 in R studio (Love et al., 2014) and defined differentially expressed genes (DEGs), when |Log_2_Fold change (FC)| ≥2 and a p-value ≤ 0.01. We evaluated a filter-bias DEGs detection by performing a linear regression between Log2FC value thresholds (1, 1.5 and 2) against the total number of DEGs detected in all thresholds. This analysis was performed for larvae, females and males separately (Figure S2b). A fit linear model was obtained across the DEGs and the thresholds using the ‘trendline’ feature of Microsoft Excel v. 2503, compilation 16.0.18623.20116, 64-bits (Microsoft Corporation 2024). A linear correlation between the total DEGs and Log_2_FC thresholds was interpreted as the lack of any bias imposed by the selected parameters, that is (1) a decrease in the total DEGs when the threshold stringency increased and (2) no major variation of the total DEGs, even with the highest threshold value. To identify DEGs associated with HW-exposure in larvae, we performed pairwise comparisons between HW-exposed samples and samples maintained under standard conditions (Table S3). For adults since the HW lasts two days we first verified any possible bias due to mosquito aging. We identified DEGs between As6 and As9 samples (List B, 99 DEGs), between As6 and As9HW (List A, 158 genes) and As9 and AsHW9 samples (List C, 113 DEGs) (Figure S1b, Table S4). We reasoned that List B contains DEGs attributable only to age differences, while Lists A and C contain DEGs that that might be affected either by both age and HW, or just one of them. If our HW regime were to induce quiescence, we would expect the transcriptome of As9HW samples to be more similar to that of As6 rather than As9 samples. However, List A (DEGs between As6 and As9HW) includes 158 DEGs, only 13 of which are shared with list B (DEGs between As6 and As9). On this basis, we conclude that it is very unlikely that our HW regime halted development. As a consequence, to understand the impact of HW on the transcriptome of adults, we focused on DEGs of List C (As9 vs As9HW) (Table S4). Shared DEGs between these lists were obtained with Venn diagrams using the online tool “Interactivenn” (Heberle et. al., 2015). The distribution and the statistical significance of DEGs exposed to a HW were shown in volcano plots, using the R package EnhancedVolcano v1.24.0 (Blighe et al., 2025). To identify genes that significantly deviate from an expected distribution, suggesting highly expressed genes, we identified outliers across each DEGs distribution in larvae, females and males, using a two-sided interquartile range test (Ramachandran &Tsokos, 2020).

### 2.8 Functional gene annotation and gene enrichment analysis

We extracted gene functions of the shared DEGs among the three above-described lists from the reference genome AalbF5, followed by manual comparison of the gene functions, as well as a GO enrichment analysis (Table S5). As shown in Figure S1b, we used only the DEGs present in List C for further analysis since they exhibited an association to HW response, suggesting thermal stress, which is in agreement with previous literature (Ware-Gilmore et al., 2023). GO functional gene assignment and gene enrichment were carried out with RStudio v.4.4.1 based on developmental stage, sex, and treatment. We performed GO functional gene assignment and gene enrichment analysis as described by Lozada-Chávez et al., (2025). Briefly, we generated an annotation database of the AalbF5 genome assembly from results of three approaches, which were merged with Blast2GO (Götz et al., 2008). The three approaches are: (1) GO annotations spanning around 72% of 20,424 protein coding genes of the AalbF5-proteome (RefSeq accession number GCF_035046485.1) obtained from NCBI RefSeq database (O’Leary et al., 2016); (2) a BLAST search of homologs of the AalbF5-proteome against the NCBI Diptera nr database v5; (3) a search of functional analogs with InterProScan v5 (Jones et al., 2014) against the protein-domain databases Pfam v33.1 (Mistry et al., 2021), ProSiteProfiles v20.2 (Sigrist et al., 2013), SUPERFAMILY v2.0 (Wilson et al., 2009) and TIGRfam v15.0 (Haft et al., 2003). From this approach, we annotated 82% of the AalbF5-proteome. Then, we performed the GO enrichment analyses using our in-house R-package org.Aalbopictus.eg.db (merged GO annotations) with clusterProfiles v4.2.2 (Wu et al., 2021). We corrected the obtained p-values (p < 0.05) for multiple tests using the Benjamini-Hochberg procedure, and we removed the redundancy of enriched GO terms for each major GO classification using the function simplify, both included in clusterProfiler.

### 2.9 Microbiota analysis

We sampled three pools of 10 L2 immediately after HW exposure and the same number of larvae maintained under standard conditions as control. We also collected eight sugar-fed females after HW exposure or, as control, eight sugar-fed females not exposed to the HW and of the same age as those HW-exposed. We sterilized all samples by rinsing them in a 2% sodium hypochlorite solution for 10 minutes, followed by surface disinfection for 5 minutes with 1x phosphate buffered saline (PBS), 70% ethanol for 2 minutes and two more rinsings with sterile water. Afterwards, we dissected the abdomens of each female. We used the Wizard® Genomic DNA Purification Kit (Promega) for DNA extraction of larval pools and single abdomens following the manufacturer’s instructions. We resuspended DNA in 20 uL nuclease-free water. Samples were analysed through next-generation sequencing (NGS) using a bacterial 16S rRNA gene target. NGS experiments were conducted by Genomix4life S.R.L. (Baronissi, Salerno, Italy). We assessed DNA quality using a NanoDrop One spectrophotometer (Thermo Scientific, Waltham, MA) and a Qubit Fluorometer 4.0 (Invitrogen, Carlsbad, CA). Library preparation targeted the hypervariable V3–V4 region of the 16S rRNA gene using the oligonucleotides 16S Amplicon PCR F (5’-TCGTCGGCAGCGTCAGATGTGTATAAGAGACAGCCTACGGGNGGCWGCAG-3’) and 16S Amplicon PCR R (5’-GTCTCGTGGGCTCGGAGATGTGTATAAGAGACAGGACTACHVGGGTATCTAATCC-3’) (Klindworth et al., 2013). We assembled each PCR reaction according to the Metagenomic Sequencing Library Preparation protocol (Illumina, San Diego, CA). A negative control was included in the workflow, consisting of all reagents used during sample processing (16S amplification and library preparation) but without template DNA, to ensure the absence of contamination. We quantified the libraries using a Qubit fluorometer (Invitrogen, Carlsbad, CA) and pooled them to achieve an equimolar concentration of each index-tagged sample, resulting in a final concentration of 4 nM, including the PhiX Control Library. The pooled samples underwent cluster generation and were sequenced on the MiSeq platform (Illumina, San Diego, CA) using a 2 × 250 paired-end format. We conducted metagenomic analysis using QIIME2 (Boleyn et al., 2019), version 2023-5.1. The 16S rDNA V3-V4 region amplicons paired-end sequence data was imported into the qiime2 program. We merged the reads using “vsearch” and quality filtered. De-noising and quality control were performed with “Deblur denoise-16S”. The qiime2 command “Features-table filter-features” was used to remove features appearing in only one sample and below 1% (Boleyn et al., 2019). Taxonomy was assigned to OTUs by using the sklearn method and aligning against the SILVA138.1 database, downloaded from the SILVA website (Quast et al., 2013). To calculate the alpha and beta diversity, we used “qiime phylogeny” and “qiime diversity core-metrics-phylogenetic” commands. The “qiime diversity alpha-rarefaction” command was used to obtain the alpha-rarefaction curves. We used the package ‘ape’ (Paradis & Schliep, 2019) in the software R (v 4.1.2; https://www.R-project.org/) and Graph Pad Prism 10.0 [Graphpad software], to plot graph and for statistical analysis, respectively.

### 2.9 Statistical analyses

We analysed fitness and physiological data using GraphPad Prism 8.0.1. We checked whether datasets conformed to a normal distribution using the Shapiro-Wilk test (Ramachandran & Tsokos, 2020). Data of HW-related mortality, hatching rate, LDT, PDT, LV, PV, sex ratio, fecundity, fertility, blood feeding rate, trypsin activity and quantification of proteins in females offered a blood meal after HW exposure were normally distributed so we used a parametric two-way ANOVA, followed by a Tukey’s multiple comparison test for post-hoc analyses (Table S6-8) (Lee & Lee, 2018). Energy reserve data did not follow normal distribution, so we assessed differences using a Kruskal-Wallis test followed by Dunn’s multiple comparison test (Table S8) (Ramachandran & Tsokos, 2020). We compared the survival of heatwave-exposed mosquitoes through a Kaplan-Meier analysis and Mantel-Cox test (Goel et al., 2010).

## 3. RESULTS

### 3.1 Eggs exposed to a HW suffer a cost in viability and hatching time

After egg exposure to our HW, we observed a significant reduction of hatching rate in both strains (Figure 2a; p < 0.05 for Cr and p<0.01 for Fo). Total hatching rate decreased from 98% (±1) to 55% (±1) in Cr and from 83% (±3) to 43% (±9) in Fo. To verify if this decrease was due to a halt in egg hatching during the HW, we followed hatching for 10 days and drew daily hatching rate curves (Figure 2b). Although the hatching rate of HW-exposed eggs started to increase two days after the HW, it remained lower than that of control mosquitoes for both strains over the course of ten days. In both strains, we observed no more than 6% hatching during the HW, while the hatching rate of controls reached daily values of 40% and 20% for Cr and Fo, respectively. Overall, these results indicate that egg exposure to extreme heat impacts egg viability and results in a delay in hatching time.

**Figure 2.**
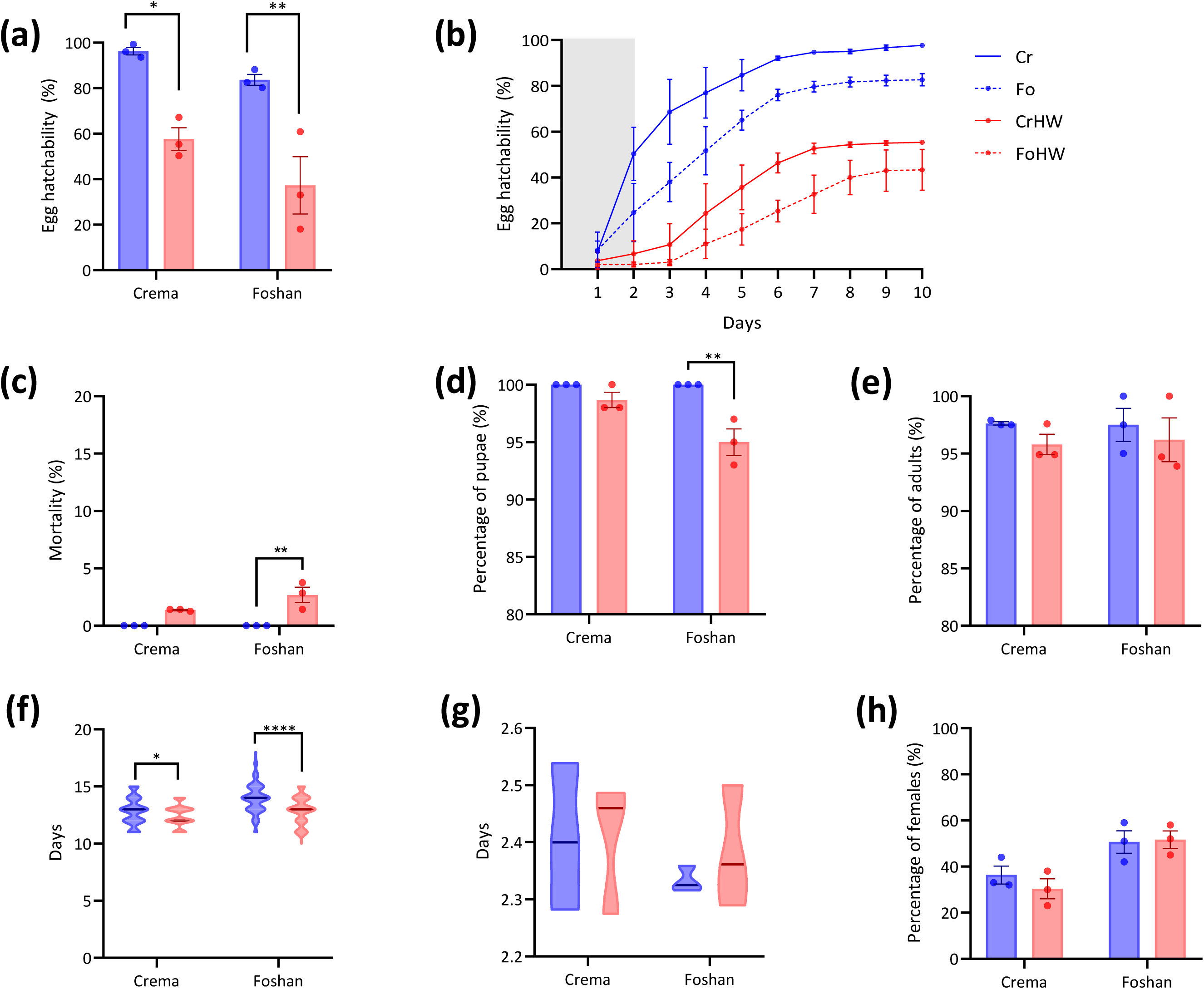
Effects of HW on eggs and juvenile stages. (a) Hatching rate. (b) Percentage of egg hatchability per day of eggs of Cr (solid line) and Fo (dotted line) mosquitoes. The grey area marks the two days of HW. (c) Mortality of larvae. (d) Larval and (e) pupal viability. (f) Larva to pupa developmental time and (g) pupal developmental time in days. (h) Percentage of emerging females (sex ratio). In all panels, red shows data for HW exposed mosquitoes and blue data of the unexposed counterparts. Statistical differences were tested through a two-way ANOVA. Error bars show the standard error of the mean (SEM). An * refers to a p-value <0.05, ** to a p-value <0.01 and **** to a p-value <0.0001.

### 3.2 Larvae are resilient to a HW

We exposed L2 to the HW, hereafter L2HW, and tracked their viability and development (Figure 1). We observed a HW-related mortality of 1.5±0.06% and 4±0.68% in Cr and Fo larvae, respectively (Figure 2c). Larval viability was significantly (p=0.0032) reduced following HW only in Fo mosquitoes (Figure 2d). LDT significantly increased in L2HW in comparison to L2 controls in both strains; LDT increased from 12.3±0.9 to 12.7±1.2 days (p = 0.047) in Cr and from 12.7±1.2 to 13.8±1.3 days in Fo (p<0.0001) (Figure 2f).We did not observe carry-over effects on PDV, adult emergence rate and sex ratio (Figure 2e,g-h).

At the transcriptome level, L2HW showed 295 DEGs with respect to L2 controls; 41 DEGs were upregulated and 254 downregulated in L2HW vs L2 (Figure 3a, Table S3). Among the genes that were highly upregulated in L2HW, we detected members of the family *protein lethal(2)essential for life* (i.e. LOC109422423, LOC115265018, LOC115265016, LOC109422406, LOC109422446 and LOC109422456) and members of the heat-shock protein (HSP) 70 family (i.e. LOC115265080, LOC109423617, LOC115266509 and LOC115265081) (Figure 3a), resulting in an enrichment for functions related to protein folding and response to heat (Table 1, Figure 3d). Among DEGs that were downregulated in L2HW, we observed an enrichment for genes associated with external encapsulating structures (i.e. LOC109414739, LOC109409014 and LOC109399534), extracellular matrix components (i.e. LOC115262565, LOC115260267 and LOC134287937) and constituents of larval cuticle (i.e. LOC109418197, LOC109413996, LOC115260267, LOC109430468) (Table 1, Figure 3d). Overall, transcriptomic results suggest that L2HW are redirecting energies to thermoregulate and pausing development, which is concordant with the observed delay in their development (Figure 2f).

**Figure 3.**
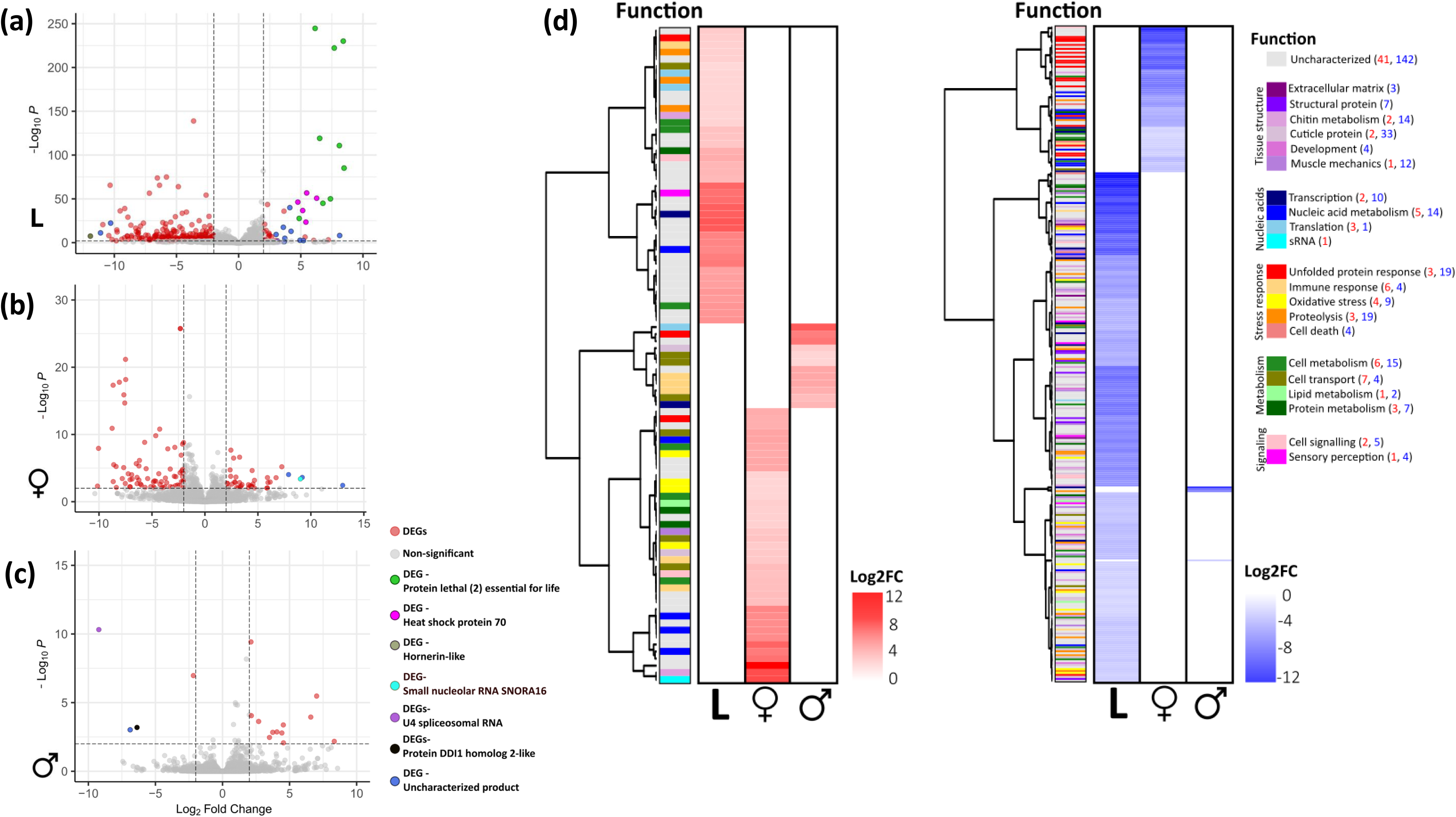
Transcriptional differences across life stages of the mosquito after a HW exposure. Volcano plots of the Differentially Expressed Genes (DEGs) in (a) larvae (L), (b) females (♀) (c) males (♂). In all cases, grey dots represent non-differentially expressed genes and red dots show DEGs for comparison between HW-exposed vs control samples. In each panel, dotted lines indicate the thresholds for the p value (<0.01) and the Log_2_FC (<|2|). Outliers DEGs are coloured based on their function, as described in the legend. (d) Heatmaps showing the upregulated (red) and downregulated (blue) genes in HW-exposed vs control L, female and male samples, along with their function. The numbers in parentheses next to the legend indicate the amount of up (red) or downregulated (blue) genes for each category.

**Table 1.** Gene Ontology enrichment of DEG after a HW exposure across life stages. * represents a set of genes with two GO terms of equal relevance

| GO | ID | Description | Regulation | Gene Ratio | Bg Ratio | p adj. | Gene ID |
| --- | --- | --- | --- | --- | --- | --- | --- |
| BP | GO:0042026 | protein refolding* | Up | 6/36 | 11/6222 | 1.82E-09 | LOC109422423, LOC115265018, LOC115265016, LOC109422406, LOC109422446, LOC109422456 |
| BP | GO:0009408 | response to heat* | Up | 6/36 | 14/6222 | 3.89E-09 | LOC109422423, LOC115265018, LOC115265016, LOC109422406, LOC109422446, LOC109422456 |
| MF | GO:0044183 | protein folding chaperone | Up | 4/59 | 24/7600 | 0.00078 | LOC115265080, LOC109423617, LOC115266509, LOC115265081 |
| BP | GO:0006950 | response to stress | Down | 8/36 | 297/6222 | 0.00636 | LOC109621883, LOC109418620 |
| CC | GO:0030312 | external encapsulating structure | Down | 15/76 | 209/5549 | 1.47E-06 | LOC109414739, LOC109409014, LOC109399534, LOC109404309, LOC109404314, LOC109404300, LOC109404318 |
| CC | GO:0031012 | extracellular matrix | Down | 15/76 | 209/5549 | 1.47E-06 | LOC115262565, LOC115260267, LOC134287937 |
| CC | GO:0005615 | extracellular space | Down | 17/76 | 367/5549 | 6.33E-05 | LOC109400176, LOC109408251, LOC115257192, LOC115257573, LOC109427011, LOC134290023, LOC109396976 |
| MF | GO:0042302 | structural constituent of cuticle | Down | 5/59 | 57/7600 | 0.00137 | LOC109419880 |
| MF | GO:0004252 | serine-type endopeptidase activity | Down | 9/59 | 253/7600 | 0.00191 | LOC109432597, LOC109419403, LOC134285492, LOC109426652, LOC109432998, LOC115253681, LOC134286581, LOC134285010, LOC109418745 |
| MF | GO:0008010 | structural constituent of chitin-based larval cuticle | Down | 4/59 | 45/7600 | 0.00321 | LOC109418197, LOC109413996, LOC115260267, LOC109430468 |
| MF | GO:0030246 | carbohydrate binding | Down | 3/59 | 47/7600 | 0.04242 | LOC134288466, LOC109418338, LOC115268558 |

B. Females
| GO | ID | Description | Regulation | Gene Ratio | Bg Ratio | p adj. | Gene ID |
| --- | --- | --- | --- | --- | --- | --- | --- |
| BP | GO:0042026 | protein refolding* | Down | 6/24 | 11/6222 | 8.71E-11 | LOC109422456, LOC109422406, LOC115265016, LOC109422423, LOC115265018, LOC109422446 |
| BP | GO:0009408 | response to heat* | Down | 6/24 | 14/6222 | 1.87E-10 | LOC109422456, LOC109422406, LOC115265016, LOC109422423, LOC115265018, LOC109422446 |
| MF | GO:0044183 | protein folding chaperone | Down | 6/41 | 24/7600 | 6.96E-08 | LOC115265081, LOC115266509, LOC115265080, LOC109423617 |
| BP | GO:0030433 | ubiquitin-dependent ERAD pathway | Down | 2/24 | 24/6222 | 0.03873 | LOC109413675, LOC109413676 |

C. Males
| GO | ID | Description | Regulation | Gene Ratio | Bg Ratio | p adj. | Gene ID |
| --- | --- | --- | --- | --- | --- | --- | --- |
| BP | GO:0006955 | immune response* | Up | 2/2 | 28/6222 | 2.79E-05 | LOC109399074, LOC134291823 |
| BP | GO:0006950 | response to stress* | Up | 2/2 | 297/6222 | 0.00227 | LOC109399074, LOC134291823 |

### 3.3 Sugar-fed males suffer a higher cost from HW exposure than females

We tested the impact of HW on 5-7-day-old sugar-fed mosquitoes, hereafter AsHW. In both Fo and Cr, we observed AsHW females suffered low mortality (3% ±0.49 in Cr and 8% ±1.19 in Fo; Figure 4a), but more than half of HW-exposed males died; percentage of male mortality was 55% ±4.60 and 76% ±16.89 in Cr and r Fo males, respectively (Figure 4b). Despite this striking sex difference in the ability to overcome HW exposure, the longevity of Cr mosquitoes surviving HW exposure did not differ drastically either between sexes or with respect to control mosquitoes (Figure 4c-d). In contrast, Fo males that had been exposed to, and had survived the HW showed a lower survival probability with respect to their control (Mantel-Cox, p=0.03). We further quantified energy reserves in both male and female AsHWs with respect to controls, named As. In accordance with longevity data, we saw that males and females differently modulate content of proteins, carbohydrates, glycogen and lipids following a HW (Figure 4e-l). We observed a significant decrease in protein content in AsHW females of both strains (p<0.0001 in Cr and p < 0.01 in Fo); in males this phenotype was observed only in Cr (p<0.01). We further saw that the content of carbohydrates and lipids did not change in males of either strain after HW, but significantly (p=0.0013) increased in Cr females (Figure 4 f,h,j,l). The opposite trend was observed in glycogen, which did not change in females, but increased in HW-exposed and surviving males.

**Figure 4.**
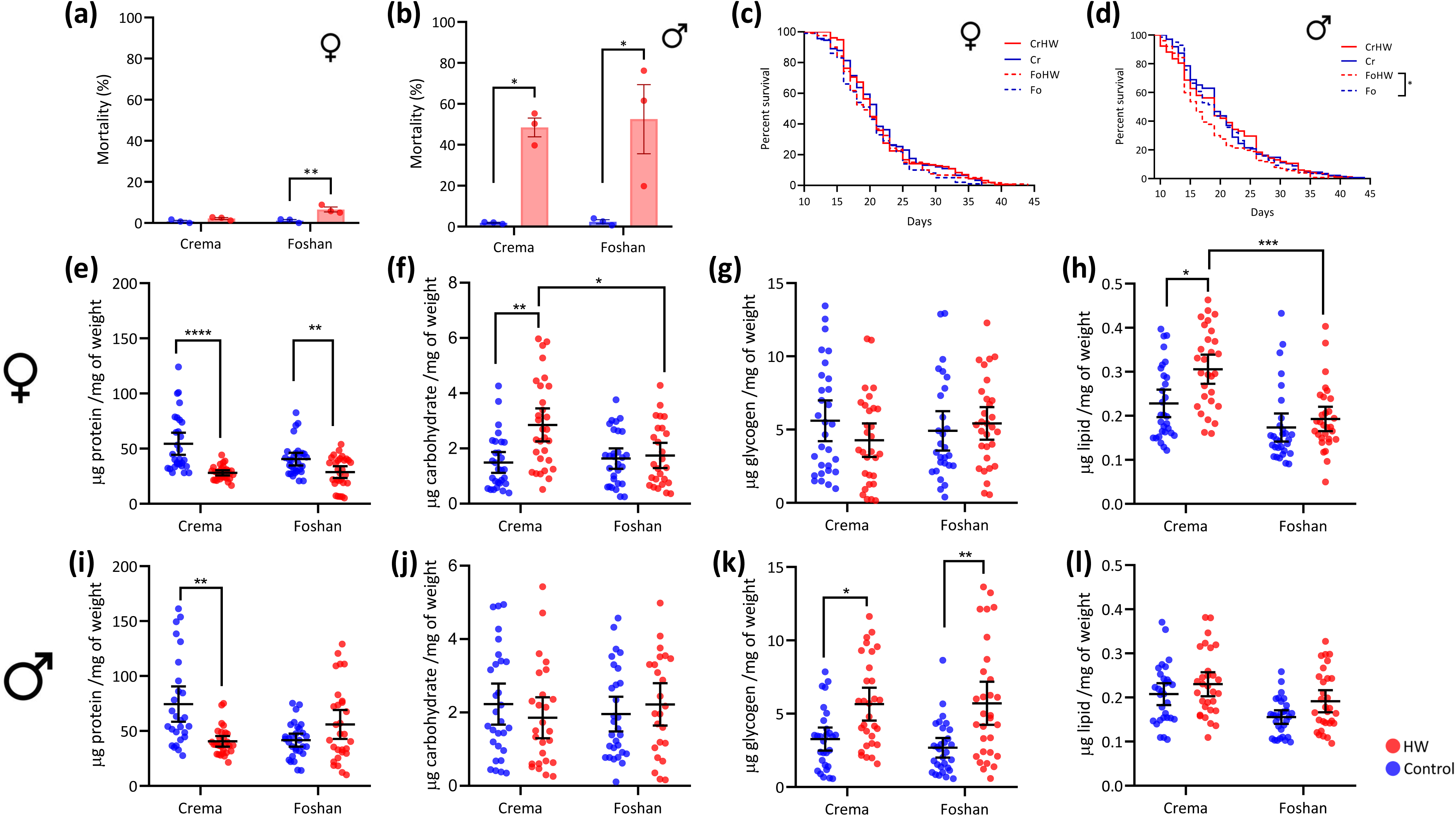
Effects of a HW in adults. Mortality of females (a) and males (b). Kaplan-Meier survival curves of adults of HW-exposed (red) and control (blue) females (c) and males (d) of Cr (solid line) or Fo (dotted line) strain. Changes in protein (e), carbohydrate (f), glycogen (g) and lipid (h) content in HW-exposed or unexposed females. Changes in protein (i), carbohydrate (j), glycogen (k) and lipid (l) content in HW-exposed and surviving males and their unexposed counterparts. In all cases, red dots show individuals that experienced a HW, whereas blue dots represent those kept at standard rearing conditions (controls). To test for statistical significance, a two-way ANOVA was used for HW-related mortality, a Mantel-Cox test for the survival curves, and a Kruskall-Wallis test for all energetical compartments. Error bars show the SEM. An * refers to a p-value <0.05, ** to a p-value <0.01, *** to a p-value <0.001 and **** to a p-value <0.0001.

At the transcriptome levels, we observed 113 DEGs, 39 upregulated and 74 downregulated in AsHW vs As females (Figure 3b, Table S4). Among upregulated genes, we detected genes related to several functions such as response to heat (i.e. LOC109422434), oxidative stress (i.e. LOC109408978, LOC109428787, LOC115260130 and LOC134288797), cell metabolism (e.g. LOC109397825 and LOC109417381) and immune response (i.e. LOC134284231 and LOC115256529). However, we could not identify any functional enrichment, probably due to the high percentage (45.1%) of uncharacterized genes (Figure 3d; Table S4). On the contrary, downregulated genes were enriched in genes associated with response to heat (i.e. LOC109422456, LOC109422406, LOC115265016, LOC109422423, LOC115265018 and LOC109422446), protein folding chaperones (i.e. LOC115265081, LOC115266509, LOC115265080 and LOC109423617) and members of the ubiquitin-dependent ERAD pathway (i.e. LOC109413675 and LOC109413676) (Table 1). In males, the number of DEGs was lower with respect to females, with 12 upregulated and 4 downregulated genes in AsHW males with respect to controls (Figure 3c, Table S4). Among upregulated genes, we observed enrichment for immunity functions (LOC109399074 and LOC134291823) and response to stress (LOC109399074 and LOC134291823) (Table 1); while downregulated genes correspond to U4 spliceosomal RNA (LOC115262321), the protein DDI1 homolog 2-like and (LOC134290813) and two uncharacterized genes (LOC134289466 and LOC115262321). Overall fitness, physiological and transcriptomic data, with the number of DEGs in males and their associated functions, support the conclusion that while females are resilient to a HW, males have limited coping mechanisms against a hot event above their thermal optimum.

### 3.4 Exposure to a HW after a blood meal results in reduced fertility

We then proceeded to explore the impact of the HW on the reproductive capacity of males and females. First, we crossed virgin males, which had been exposed and survived the HW, to unexposed virgin females and observed no changes in fecundity and fertility (Figure 5a-c). Next, we studied the reproductive capacity of females and tested the impact of exposing females to a HW either before (AHWb) or after (AbHW) a blood-meal. In AHWb, we observed a reduction in blood-feeding rate in Cr (p=0.0195; Fig. 5d), but no significant differences in fecundity and fertility (Figure 5e,f). On the contrary, in both strains, we saw a significant reduction of fertility in AbHW (p<0.001 in Cr and p=0.0061 in Fo; Figure 5e,f). Neither AHWb nor AbHW showed changes in sterility rate (Figure 5g).

**Figure 5.**
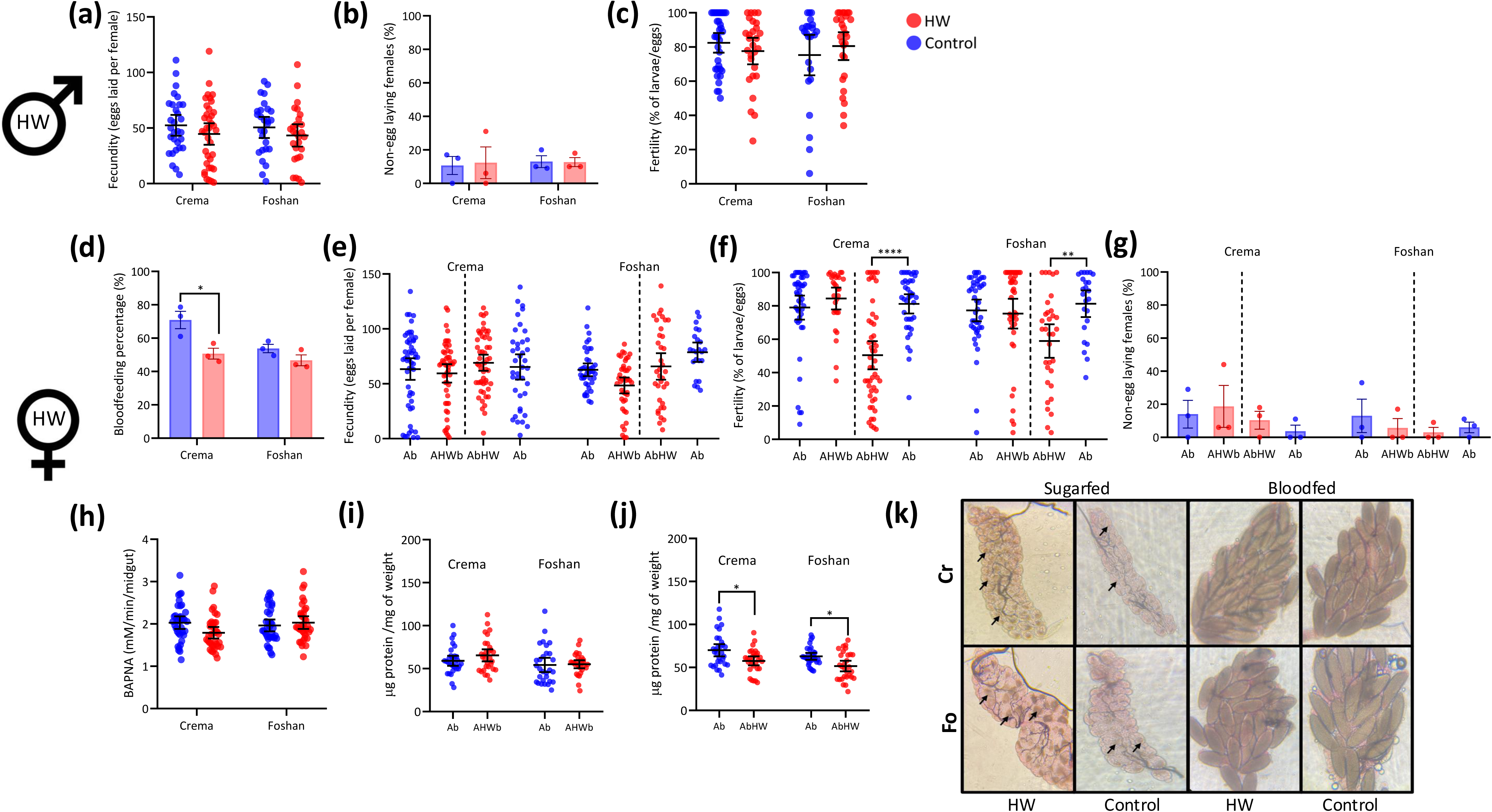
Reproductive traits of HW survivors. (a) Fecundity, (b) percentage of sterile females and (c) fertility in virgin females mated with HW-exposed males. (d) Bloodfeeding rate of AHWb females. (e) Fecundity, (f) fertility and (g) percentage of sterile females of AHWb and AbHW females. (h) Trypsin-like activity of AHWb 24h after the bloodmeal. (i) Protein content of AHWb females 3 days post blood meal. (j) Protein content of AbHW immediately after the HW. (k) Ovarian micrographs stained with neutral red of AsHW and AbHW females. Arrows indicate the lysosomes in the follicles. In all cases, red and blue dots represent data from HW-exposed or control (Ab) mosquitoes. Control mosquitoes were sampled to be of the same age of HW exposed mosquitoes. Statistical significance was tested using a two-way ANOVA. Error bars show the SEM. An * refers to a p-value <0.05, ** to a p-value <0.01, *** to a p-value <0.001 and **** to a p-value <0.0001.

Given that we had observed a reduction of protein in AsHW females (Figure 4e), but AHWb females showed no reduction in their reproductive output (Figure 5ef), we hypothesized nutrients from the blood-meal compensate for HW-associated protein reduction. To test this hypothesis, we measured trypsin-like activity 24 hours post blood-meal (Figure 5h) and protein content of AHWb females 3 days after the blood meal. We observed no difference in trypsin-like activity (Figure 5h) nor protein content (Figure 5i) between AHWb and their controls, confirming that nutrients from a blood-meal can compensate for HW-associated protein reduction. We further measured protein content of AbHW immediately after the HW and checked the state of ovaries of AbHW and AsHW females since limited availability of proteins can result in ovarian resorption (Clifton & Noriega, 2011). In both strains, we saw a decrease in protein content in AbHW with respect to Ab females (p>0.05; Figure 5j), but this reduction was not as drastic with respect to what we had observed in AsHW females (Figure 4e). Additionally, we did not see differences in follicle resorption in either AsHW or AbHW versus control females (Figure 5k). We observed that eggs of AsHW females tend to have larger lysosomes than those observed in As females (Figure S3), suggesting no resorptions, but alterations of egg physiology.

### 3.5 HW alters larval but not female composition of the microbiota

Finally, we checked whether HW exposure impacts the composition of the microbiota of L2 and As females. In larvae, we identified a total of 29 bacterial taxa (Figure 6a). We observed that the microbiota of one L2HW sample of the Cr strain consisted for > 90% of *Stenotrophomonas*, a taxon we did not observe in any other sample. To avoid any possible bias, we excluded data from this sample in further analyses. In general, we observed a great variability in the microbiota composition of L2HW samples with a decrease in *Microbacterium* and *Methylobacterium* and an increase in *Sphingomonas* and *Acinetobacter* in both strains. We also saw strain-specific blooming of different bacterial taxa, but no changes in *Wolbachia* abundance between L2 vs L2HW samples. Briefly, *Microbacterium* decreased from 32.8 – 47.3% to 3.3 – 9.2% in L2 vs L2HW Cr samples and from 41.1 – 55.0 % to 7.8 – 21.2% in L2 vs L2HW Fo samples (Table S9); *Methylobacterium* decreased from 1.6 – 23.6% to 0.2 – 13.1% in L2 vs L2HW Cr samples and from 7.8 – 21.2 % to 2.0 – 7.0 % in L2 vs L2HW Fo samples. On the opposite, *Sphingomonas* increased from 0.4 – 1.7 % to 21.1 – 32.6 % in L2 vs L2HW Cr samples and from 0.3 – 3.1 % to 5.6 – 24.2% in L2 vs L2HW Fo samples and *Acinetobacter* increased from 0.1% to 1.8 – 25.6% in L2 vs L2HW Cr samples and from 0.2 –0.9 % to 1.5 – 46.6% in L2 vs L2HW Fo samples. Strain-specific blooming was observed for *Delftia* and *Chryseobacterium. Delftia* increased from 0.0–0.3% to 0.5–2.4% in L2 vs L2HW Cr samples; while *Chryseobacterium* increased from 0.1–0.5% to 3.2–17.7% in L2 vs L2HW Fo samples (Table S9). *Wolbachia* abundance ranged from 6.7–23.3% to 4.3–23.4% in L2 vs. L2HW Cr samples and from 5.5–13.8% to 5–18.9% in L2 vs. L2HW Fo samples. These results were reflected in the beta-diversity analyses, which showed larval samples clustering based on treatment, but not strain and L2 samples having a more homogenous microbiota composition than L2HW ones (Figure 6b). A similar trend as described for beta-diversity analyses was observed in the Weighted UniFrac analysis (Fig. 6c).

**Figure 6.**
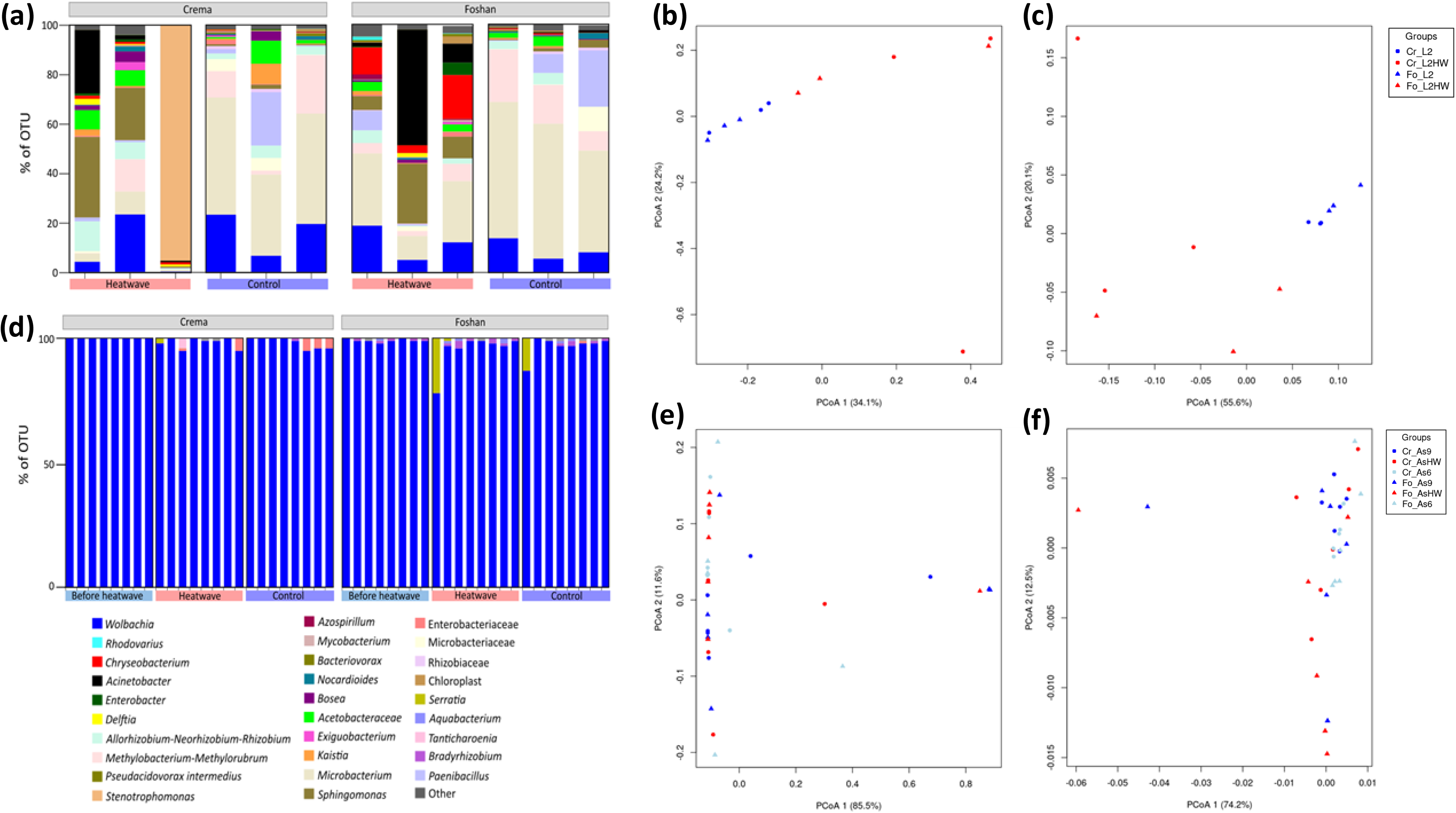
Changes in bacterial composition after a HW. (a) Barplot of bacterial taxa in L2 and L2HW. Brays-Curtis and Weighted-UniFrac analyses of larvae (b, c), respectively. (d) Barplot of bacterial taxa of As and AsHW samples, with their respective Bray-Curtis (e) and Weighted-UniFrac analyses (f). In each panel, data for Fo an Cr mosquitoes are in triangle or circle, respectively. Red shows data from HW-exposed samples, blue to data of control samples. For females, we collected unexposed control samples 6 (light blue) and 9 (dark blue) days post-emergence to match the age before and after HW-exposed samples before and after HW exposure.

The microbiota of As samples was dominated by *Wolbachia*, independently from treatment and strain (Fig. 6d). *Wolbachia* ranged from 95.6%-100% to 77.9%-100% in Cr vs Fo samples, respectively (Table S9). As shown by the beta-diversity analyses, the composition, and the relative abundance of bacterial taxa, was overall more homogenous across adults than larval samples and the microbiota of adult samples was also less impacted by HW exposure (Figure 6e-f).

## 4. DISCUSSION

To cope with the thermal stress resulting from exposure to extreme temperatures as during a HW, holometabolous insects may use stage-specific mechanisms that define the thermal sensibilities of each life instar (González-Tokman et al., 2020). Here, we show the phenotypic, physiological and transcriptional changes occurring across the development of *Ae. albopictus* when exposed to an ecologically relevant HW, in which the nightly drop of 11°C from 37° during the day served as a recovery period (Bai et al., 2019; Ma et al., 2021; Zhu et al., 2019).

We observed that larvae and females are HW resilient, while eggs and males suffer extensive fitness and physiological costs. The egg hatching rate that we observed following HW is consistent with what reported when hatching *Aedes aegypti* eggs at constant 37°C (Pekľanská et al., 2025). This indicates that hatching occurred both during the hot and the recovery periods of our HW regime. However, the fact that the overall hatching rate of HW exposed eggs was lower than that of control eggs suggests embryo mortality due to elevated temperature. Aquatic larvae are expected to be sensitive to heat (González-Tokman et al., 2020), but our results showed that mosquito larvae can overcome HW exposure by delaying their development. The hypothesis of larval developmental delay is supported by our transcriptional data, which showed lower expression of growth-related genes (i.e. *apnoia* and *agrin*), multiple cuticle and chitin-related genes, genes encoding for muscle-related proteins (e.g. *titin*) and genes involved in cornification (i.e. *hornerin-like* and *loricrin*) in L2HW with respect to L2 larvae. In *Ae. aegypti,* heat response of larvae is very dynamic, with a higher HSP expression than in other life stages (Zhao et al., 2010). We also found the highest number of HSPs was elicited in larvae than adults following HW exposure. Upregulated HSPs included members of the *protein lethal(2)essential for life*, which belong to the family of *hsp20*, and members from the *hsp70* family. Both families of chaperones are known to contribute to thermal tolerance in insects (Mack & Attardo, 2024).

At the adult stage, we observed extensive sex-differences in response to HW exposure. We saw that males surviving HW-exposure accumulated glycogen. Glycogen is mainly used to fuel short-term flight in insects; its increase suggests males reduce movement during HW (Arrese & Soulages, 2010), which can limit their range shifts to more favourable microclimates. In accordance with this hypothesis, we saw more than 50% male mortality following HW exposure supporting the conclusion that HWs can be a strong selective pressure for males, ultimately impacting mosquito population size. At the transcriptome level, males surviving HW exposure upregulated mostly immunity genes such as the antimicrobial peptides defensin A, B and C. Defensins are elicited upon tissue damage to prevent infections (Ueda et al., 2005), supporting the hypothesis of heat-induced tissue damage in males. Differently than males, females showed less than 20% HW-related mortality and extensive transcriptional responses. We found several members of the *hsp70* (i.e*. heat shock protein 70 B2, heat shock protein A1-like, heat shock protein 70 A1*) and *hsp20 (*i.e. *protein lethal(2)essential for life, protein lethal(2)essential for life-like)* families, which are upregulated in L2HW, being downregulated in HW-exposed females, further highlighting the stage-specificity of these proteins. Expression of HSPs is dynamic and dependent both on the length and intensity of the thermal challenge (González-Tokman et al., 2020) because a high expression of HSP*s* is costly and may interfere with several cellular processes resulting in toxicity (Feder & Hofmann, 1999). Downregulation of *hsp70* and *protein lethal(2)essential for life* in females following HW exposure may impact their vector competence as the replication of both Zika and dengue viruses (DENV) is positively regulated by *hsp70* (Taguwa et al., 2015, 2019). While, we saw only one member of the *hsp20*, a *protein lethal(2)essential for life* (i.e. LOC109422434) being upregulated in AsHW females, we found several genes involved in oxidative stress (ie. *Senecionine N-oxygenase*, *farnesol dehydrogenase-like, NADH-ubiquinone oxidoreductase chain 2-like, estradiol 17-beta-dehydrogenase 11-like*) and genes involved in cell metabolism (i.e. *alkaline phosphatase 4-like, regucalcin, max-like protein_X*) being highly (>2 LogFC) upregulated in AsHW, which agrees with the observed higher abundance in lipids and carbohydrates. These results suggest that females can tolerate a HW by increasing their sugar and lipid metabolism resulting in production of free radicals, which require activation of detoxification mechanisms (González-Tokman et al., 2020).

Results from investigations testing the impact of hot treatments designed with no recovery periods show extensive negative effects on the reproductive output of various insect species including *Dr. melanogaster* and other Drosophilidae, the red flour beetle *Tribolium castaneum*, the bed bug *Cimex lectularius*, the parassitoid *Aphidius avenae* and the oriental fruit moth *Graphilita molesta* through alterations of insect behaviours and gamete production and viability (David et al., 2005; Porcelli et al., 2017; Roux et al., 2010; Rukke et al., 2015, 2018; Sales et al., 2018; Zheng et al., 2017). We saw a reduction of female fertility when HW exposure occurred immediately after the blood-meal, but we did not see a reduction in either fecundity or fertility in HW-exposed males or females exposed to HW before the blood-meal. Biological effects of a hot event depend on a subtle balance between heat stress related injuries and the possibility to recover during fluctuations to milder temperatures (Ma et al., 2021). Thus, the nightly recovery period of our HW regime could have buffered the heat-related injuries from exposure to 37°C, a temperature close to the *Ae. albopictus* physiological threshold (Carlassara et al., 2024). Additionally, we exposed to the HW sexually mature mosquitoes, which, in the case of males, were allowed to mate for five days after HW exposure. We cannot exclude that potential negative HW impacts were masked by a compensatory effect in the days following the HW as male insect fertility appears to be mostly impacted when the hot event occurs before sexual maturation (Sales et al., 2021). While we recognize that additional studies mostly exposing males at emergence and restricting their mating window, in addition to experiments testing alternative HW regimes should be explored to further understand the biological impact of hot events on *Ae. albopictus*, we highlight the ecological relevance of our results. Italian meteorological records show the frequent occurrence of hot events, mostly consisting of two to three days with a thermal maximum in the range of 37-38°C followed by 24-27°C at night in the summer months for the past decade (Time and Date, www.timeanddate.com/weather/italy/rome/historic?month=8&year=2024). These thermal conditions are very similar to our HW, which was chosen to mimic a hot event recorded in Rome in August 2021 (World-Weather, https://world-weather.info/forecast/italy/rome/august-2021/).

Additionally, despite Fo and Cr mosquitoes being genetically different and having a different thermal optimum (Carlassara et al., 2024), their developmental-specific responses to our HW regime were mostly concordant, allowing for generalization. Our findings have thus important implications for predicting mosquito population dynamics under current global climate changes and organize control strategies. Our results indicate that control of juvenile stages would be effective through the summer season as larvae appear to be heat resilient and their development may lengthen in response to hot events. At the same time, the observation of a decrease in *Microbacterium* in L2HW provide indication on insecticide choices as *Microbacterium* is among the insect gut bacteria shown to be able to degrade pyrethroid and organophosphate insecticides (Malook et al., 2025). Our results on the vulnerability of males to a HW highlight the importance of programming releases of genetically-modified, or radiation-sterilised, males in relation to weather forecast to possibly avoid hot events that could decrease release efficacy. Our data on female resilience to HWs and the absence of HW-dependent fluctuations of *Wolbachia* in AsHW support the sustainability of *Wolbachia*-based control strategies also during the occurrence of thermal fluctuations to high temperatures in presence of a nightly recovery period.

## ACKNOWLEDGEMENTS

We are grateful to all members of the Bonizzoni’s lab for fruitful discussion.

## AUTHOR CONTRIBUTION

C. Alfaro performed biological experiments, contributed to bioinformatic analysis of transcriptomic data, analyzed the data and wrote the manuscript; A. N. Lozada-Chávez contributed to bioinformatic analyses of transcriptomic data, statistical analyses of the data and analyzed the data; A. Khorramnejad contributed to energy reserve experiments and analyzed the data; A. Kropf contributed to overall data analysis. P.L. Catapano performed bioinformatic analyses of microbiota data A. Cappelli analysed the microbiota data; C. Damiani analysed the microbiota data; G. Favia contributed to microbiota analysis and contributed with funding; M. Bonizzoni conceived and supervised the study, obtained funding and wrote the manuscript with feedback from all authors.

## DATA AVAILABILITY STATEMENT

Whole Genome Sequencing data that support the findings of this study are openly available in NIH SRA BioProject ID number PRJNA1272515. All raw scaled reads data counts per sample generated by our research, as well as the command lined used to run nf-core/RNAseq and a custom R script used to estimate DEGs using DESeq2 for the transcriptomic analysis were deposited in the publicly accessible repository https://github.com/naborlozada/Alfaro_et_al_2025_Aalbo_heatwaves.

## SUPPLEMENTARY MATERIAL

### Supplementary figures

**Figure S1.** Transcriptome controls for considering the 2-day time effect of the HW in adults. **(a)** Simulated heatwave conditions and sampling points for the transcriptomic analyses of adults. The blue line on the graph shows the standard rearing conditions, and the red line represents the HW conditions. Females and males were exposed to HW at 6 days old and collected on day 9 (AsHW, red) together with their unexposed counterparts (As9, blue). To address a potential two-day time effect, we further sampled females and males on day 6 post emergence (As6, light blue). (b) We obtained the list of DEGs between As6 vs AsHW (red, A), As6 vs As9 (blue, B) and As9 vs AsHW (dark red, C). To account for the “two-days time effect”, all genes of List A that are shared with List B were removed, and the remaining DEGs of List A were compared to those of List C. We selected the List C for all the rest of the transcriptional analyses performed.

**Figure S2.** Genes that are expressed across life-stages of *Ae*. *albopictus* after experiencing a HW. (a) Principal component analysis using the normalized counts of all samples. Triangles represent larvae, dots females and square males. Red and blue refer to samples exposed to a HW or kept at standard rearing conditions, respectively. (b) Linear regressions of DEGs lists when using Log_2_FC values of 1, 1.5 and 2, of larvae, females and males. For adults, the linear regressions are plotted for List A (As6 vs AsHW9, light blue), List B (As6 vs As9, blue) and List C (As9 vs AsHW9, red) of DEGs. The correlation coefficient is shown next to each line.

**Figure S3.** Assessment of ovarian physiology after a HW. Ovaries stained with neutral red of sugarfed and bloodfed females of Cr and Fo strain after the HW. The arrows indicate lysosomes present in the ovarian follicles.

### Supplementary tables

**Table S1. Scaled gene counts from individuals used for transcriptomic analyses**

**Table S2. Total number and percentages of expressed genes across life stages and treatments of *Ae. Albopictus***

**Table S3. Larvae DEGs after a HW exposure**

**Table S4. Females’ and males’ DEG Lists A, B and C.** (a) Females’ list A (As6vsAsHW9), list B (As6 vs As9) and list C (AsHW9). (b) Males’ list A (As6vsAsHW9), list B (As6 vs As9) and list C (AsHW9).

**Table S5. GO enrichment of DEGs using different controls for considering 2 days time effect**.

**Table S6. Statistic tests for significant differences in eggs and larvae fitness traits.** (a) Egg hatching rate. (b) Larvae mortality. (c) Larval viability. (d) Pupal viability. (e) Larval developmental time. (f) Pupal developmental time. (g) Sex ratio.

**Table S7. Statistic tests for significant differences in reproductive traits**. (a) Fecundity and (a) fertility of HW-exposed males. Non-egg-laying females (c) and non-fertile females (d) crossed with HW-exposed males. (e) Bloodfeeding rate of AsHWb females. (g) Fecundity and (h) fertility of AHWb females. Non-egg laying (i) and non-fertile (j) AHWb. Fecundity (k) and fertility (l) of AbHW females. Non-egg laying (m) and non-fertile (n) AbHW females.

**Table S8. Statistic tests for significant differences in adult mortality and energy reserves**. Female (a) and male (b) mortality. (c) Protein content. (d) Carbohydrate content. (e) Glycogen content. (f) Lipid content. (g) Protein compensation of AsHW females three days after bloodmeal. (h) Protein content of AbHW females.

**Table S9. Percentage of bacterial OTUs in heatwave exposed and non-exposed larvae and adults.** (a) Larvae pool samples. Female individuals of Cr (b) and Fo (c) strain.

